# Deep Learning Based Proarrhythmia Analysis Using Field Potentials Recorded from Human Pluripotent Stem Cells Derived Cardiomyocytes

**DOI:** 10.1101/244442

**Authors:** Zeinab Golgooni, Sara Mirsadeghi, Mahdieh Soleymani Baghshah, Pedram Ataee, Hossein Baharvand, Sara Pahlavan, Hamid R. Rabiee

## Abstract

**Aim:** An early characterization of drug-induced cardiotoxicity may be possible by combining comprehensive *in vitro* pro-arrhythmia assay and deep learning techniques. The goal of this study was to develop a deep learning method to automatically detect irregular beating rhythm as well as abnormal waveforms of field potentials in an *in vitro* cardiotoxicity assay using human pluripotent stem cell (hPSC) derived cardiomyocytes and multi-electrode array (MEA) system.

**Methods and Results:** We included field potential waveforms from 380 experiments which obtained by application of some cardioactive drugs on healthy and/or patient-specific induced pluripotent stem cells derived cardiomyocytes (iPSC-CM). We employed convolutional and recurrent neural networks, in order to develop a new method for automatic classification of field potential recordings without using any hand-engineered features. In the proposed method, a preparation phase was initially applied to split 60-second long recordings into a series of 5-second long windows. Thereafter, the classification phase comprising of two main steps was designed. In the first step, 5-second long windows were classified using a designated convolutional neural network (CNN). In the second step, the results of 5-second long window assessments were used as the input sequence to a recurrent neural network (RNN). The output was then compared to electrophysiologist-level arrhythmia (irregularity or abnormal waveforms) detection, resulting in 0.84 accuracy, 0.84 sensitivity, 0.85 specificity, and 0.88 precision.

**Conclusion:** A novel deep learning approach based on a two-step CNN-RNN method can be used for automated analysis of “irregularity or abnormal waveforms” in an *in vitro* model of cardiotoxicity experiments.

## Introduction

Cardiotoxicity is one of the major reasons for drug attrition from the market which may impose tremendous costs to pharmaceutical companies ^1^. Comprehensive *in vitro* pro-arrhythmia assay (CiPA) using the IPSC-CM/MEA system have been proposed to be a robust, efficient, and sensitive platform for cardiotoxicity screenings ^2-8^. While industry standard assays are based on using immortalized cell lines or animal models, CiPA takes the advantage of cardiomyocytes obtained from cardiogenic differentiation of hPSCs, literally representing the most similar physiology to human heart ^9^. Therefore, this platform may provide an advanced complementary method with great potential for reducing the costs of drug development and cardiotoxicity-related drug attrition.

Field potentials recorded from iPSC-CMs using MEA system is well correlated to action potentials recorded from single cardiomyocytes and electrocardiogram (ECG) recorded from the whole heart ^10^. Furthermore, IPSC-CM/MEA platform could reliably demonstrate the cardiotoxicity of drugs which are known to be arrhythmogenic, according to the previous experiments ^11-13^. Based on these findings, efforts have been made to generate a quantitative system in order to predict drugs with pro-arrhythmic risk ^11, 14, 15^.

By developing diverse experimental platforms and their quantitative readouts, scientists face large datasets which required extensive human resource and time to be analyzed and interpreted. Last few decades have been the golden age for the development of automated tasks to help experts and improve performance in the field of medical data analysis ^16-20^.

Especially, recent studies have proposed deep learning methods to overcome drawbacks of employing hand-engineering features to describe data characteristics. In fact, the deep learning methods do not explicitly require any type of feature design by human experts, and instead, the features are implicitly learned from data through an automatic learning procedure ^21^. More recently, a few studies have investigated the task of analysis and classification of electrocardiogram (ECG) signals with deep learning methods ^22, 23^. Rajpurkar et al. ^22, 23^ introduced a deep learning model to classify ECG samples and could exceed the average cardiologists performance in both sensitivity and precision measures ^23^. In another study, a personalized monitoring and warning system based on deep learning methods was designed for cardiac arrhythmia prediction ^22^.

Despite the development of new methods for automation of arrhythmia detection on surface ECG, the interpretation of data obtained from IPSC-CM/MEA system and the translation of findings for risk assessment is still in its infancy. Advanced machine learning methods can play an important role on creating computer-aided diagnosis (CAD) tools for IPSC-CM/MEA data analysis. In this study, we hypothesized that the deep learning approach can be exploited to create a model for irregularity or abnormal waveform detection in the field potentials recorded from healthy and/or patient-specific iPSC-CMs, which have been subjected to some cardioactive drugs. Our model is comprised of two different deep learning models, namely CNN ^24^ and RNN ^25^. Our results showed high performance of the proposed method for irregularity or abnormal waveform detection.

## Methods

### Data set creation

#### Generation of iPSC-CMs

In order to induce CMs differentiation of human iPSCs in a suspension culture system, the 5-day-old iPSC size-controlled spheroids (average size: 175 ± 25 µm) were cultured for 24 hours in differentiation medium (RPMI 1640 medium; Gibco) supplemented with 2% B-27 supplement without vitamin A (12587-010; Gibco), 2 mM L-glutamine (25030-024; Gibco), 0.1 mM β-mercaptoethanol (M7522; Sigma-Aldrich) and 1% nonessential amino acids (11140-035; Gibco) containing 12 µM of CHIR99021 (a small molecule activating canonical Wnt/β-catenin pathway) (CHIR; 041-0004; Stemgent). The spheroids were subsequently washed with Dulbecco’s phosphate-buffered saline (DPBS) and maintained in fresh differentiation medium without small molecule (SM) for one day. Thereafter, the medium was exchanged for new differentiation medium that contained 5 µM IWP2 (3533; Tocris Bioscience) as a Wnt antagonist, 5 µM SB431542 (S4317; Sigma-Aldrich) as an inhibitor of transforming growth factor-β superfamily type I activin receptor-like kinase receptors, and 5 µM purmorphamine (Pur; 04-0009; Stemgent) as a sonic hedgehog (SHH) agonist. The spheroids were cultured for 2 days in this medium, after which they were washed with DPBS and further cultured in a fresh differentiation medium without SMs. This medium was refreshed every 2–3 days.

#### Application of cardioactive drugs

We used a multi-electrode array (MEA) data acquisition system (Multichannel Systems) for extracellular field potential recordings. The MEA plates contained a matrix of 60 titanium nitride electrodes (30 µm) with an inter-electrode distance of 200 µm. MEA plates were sterilized with 70 % ethanol solution and hydrophilized with fetal bovine serum for 30 minutes, washed with sterile water, and then coated with an ECM gel from Engelbreth-Holm-Swarm mouse sarcoma (catalog no. E1270; Sigma-Aldrich). Beating spheroids were plated in differentiation medium on the middle of sterilized MEA plates for at least 48 hours. On the day of the experiment, the MEA plates were connected to a head stage amplifier. The recordings were performed for 60 seconds at baseline and 5 minutes after drug application. Verapamil hydrochloride (Sigma-Aldrich), Nifedipine (Sigma-Aldrich), Propranolol hydrochloride (Sigma-Aldrich) and Sotalol hydrochloride (Sigma-Aldrich) were used. All recordings were performed at 37°C. The extracellular field potentials were sampled at 2 KHz in Cardio2D software (V 2.6.2; Multichannel Systems) and saved for further analysis.

### Data analysis

A novel approach for automated assessment of irregularity and/or abnormal waveforms in the field potentials recorded from iPSC-CMs was proposed. The recorded field potentials were classified into normal or arrhythmic by utilizing a specific deep learning approach.

#### Data preparation

To prepare input signals from field potential recordings of iPSC-CMs, preprocessing (denoising and splitting) steps were required. First, each raw signal was normalized and filtered using a moving average filter to reduce noise. Then, the preprocessed 60-second long signals were divided into random 5-second windows, each started at a particular R-peak and continued to the fixed length of five seconds (Figure 1).

**Figure 1.**
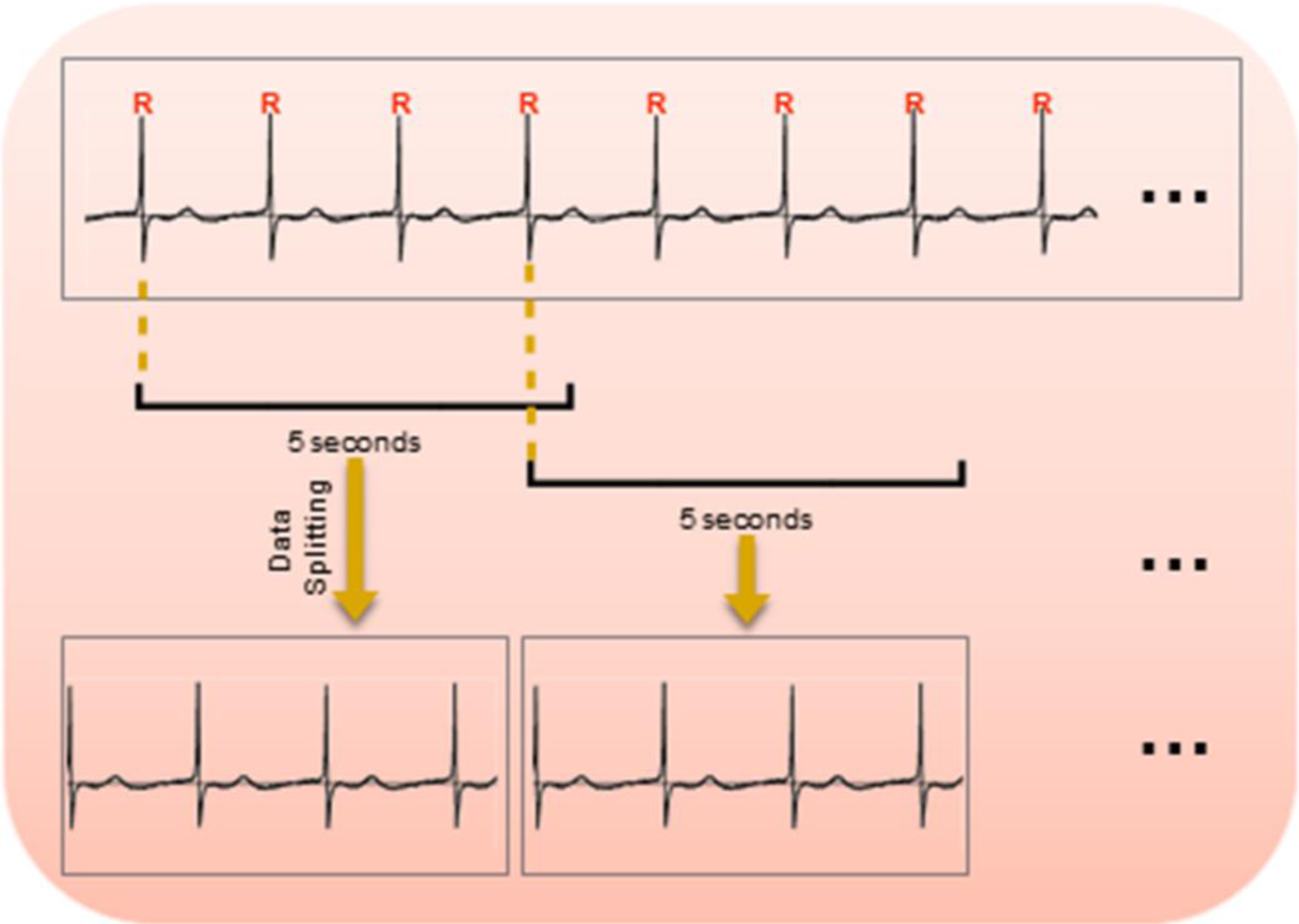
Splitting a 60-second long recording into a sequence of 5-second long windows. To split a record, R peaks are detected all over the recording. Each 5-second long window starts from a particular R point and continues to the fixed length of five seconds.

#### Classification model

After the preparation phase, the preprocessed input signals were classified into normal or arrhythmic by using a two-step deep learning architecture (Figure 2). In the first step, we designed a specific 1-D CNN architecture which used the preprocessed signals as its input without employing any hand-engineered feature. The input layer size was 5000, which was obtained from 5-second long signals at 1 KHz sampling frequency. Our network contained three layers of convolution, each had 16 filters with the kernel size of 80. Moreover, batch normalization ^26^ was adopted right after each convolution and before the activation function that was chosen to be Scaled Exponential Linear Unit (SELU) ^27^. We also used dropout ^28^ at each convolution layer to improve the generalization capability of the proposed algorithm. The network ended with one fully-connected layer and the sigmoid activation function that produced one output specifying the probability of abnormality for the corresponding window.

**Figure 2.**
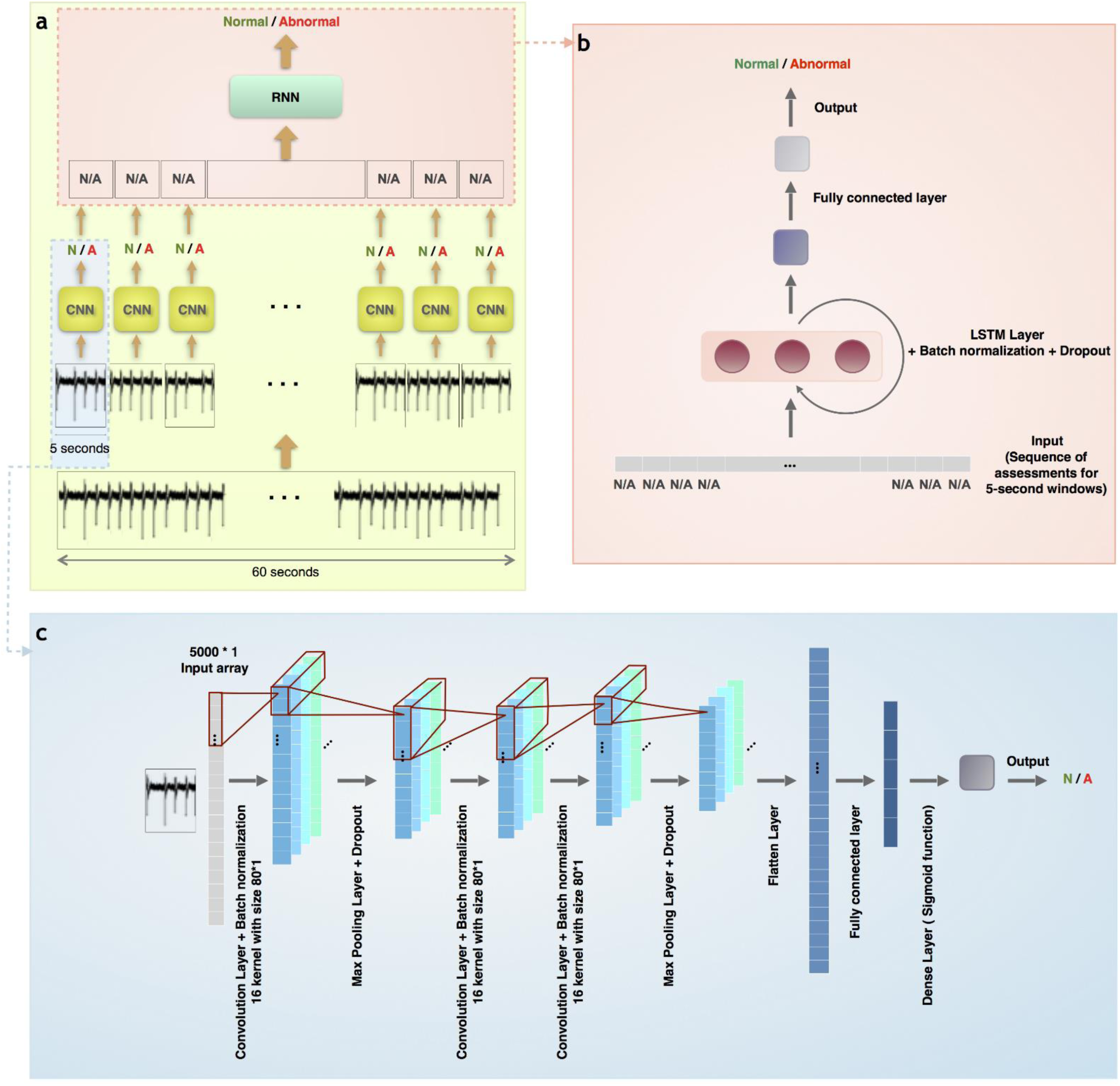
Representation of the two-step deep learning-based classification model. a) The general architecture of the proposed method: To classify a field potential recording of iPSC-CMs, a primary denoising followed by splitting of the 60-second long recording into the set of 5-second signals is applied. Each 5-second-long window is fed into the trained CNN (the details of the CNN architecture was shown in (b)) which produces output results for that window. Then, the sequence of CNN outputs is used as the input for the trained RNN (its components were presented in (c)). The RNN output determined our final result given the decisions of CNN on the 5-second signals, classifying the 60-second long filed potential recording as normal or arrhythmic. Abbreviations: N; Normal and A; Arrhythmic. b) The CNN architecture that takes 5-second long window as input and provides assessment of that window (probability of being normal or arrhythmic for 5-second long window). c) The RNN architecture that takes the sequence of 5-second long windows assessments as the input and provide the final result (probability of being normal or abnormal for 60-second long recording).

The second step was dedicated to classification of the whole field potential recording, based on the sequence of evaluation results obtained on the 5-second long windows. For this purpose, the well-known Long Short Term Memory (LSTM), ^29^ as an RNN architecture, was employed. The designed network contained one recurrent layer with three neurons and one fully-connected layer with the sigmoid activation function. In addition, batch normalization and dropout were applied in the recurrent layer.

To train the proposed model composed of the CNN and RNN networks, we used our previously generated dataset which includes both normal and arrhythmic field potential recordings of iPSC-CMs that was previously labeled by electrophysiology experts.

## Results

In this study, we developed a new classification model to automatically identify normal and abnormal field potential waveforms of human cardiomyocytes differentiated from iPSCs of healthy or patient individuals as well as their cardioactive drugs experiments (Figure 3). The healthy and patient-specific iPSC-CMs were generated using a small molecule-based differentiation protocol ^30-32^. The spheroids of 30-day old iPSC-CMs were plated on the electrodes of a multielectrode array system and their filed potential recordings were performed (Supplementary movie 1) ^30, 32^. The field potentials recorded from multi-cellular spheroids of iPSC-CMs resembled the surface ECG recorded from the whole heart (Supplementary Figure 1) and provided valuable information on the excitability and functionality of differentiated cardiomyocytes. The field potential waveform included data on the duration of a beating cycle (R-R interval), the field potential duration (FPD) which corresponds to repolarization phase of the action potential and QT interval of ECG, and the field potential rhythmicity (Supplementary Figure 2). We also applied some known cardioactive drugs with acute effects on healthy and patient-specific iPSC-CMs in order to experimentally mimic arrythmogenesis and to provide raw data for the development of classifier. Therefore, we are not going to discuss the drug effects in this paper. These experiments helped the formation of a large dataset with valuable information but time-consuming and difficult to be manually analyzed. Furthermore, the *in vitro* cardiotoxicity assay using iPSC-CMs and MEA system is a novel approach which lacks sufficient and appropriate analysis software. Although efforts have been made to develop analysis tools ^14, 15^, to the best of our knowledge, there has not been any published results on the automatic classification of field potential recordings from iPSC-CMs into normal or arrhythmic.

**Figure 3.**
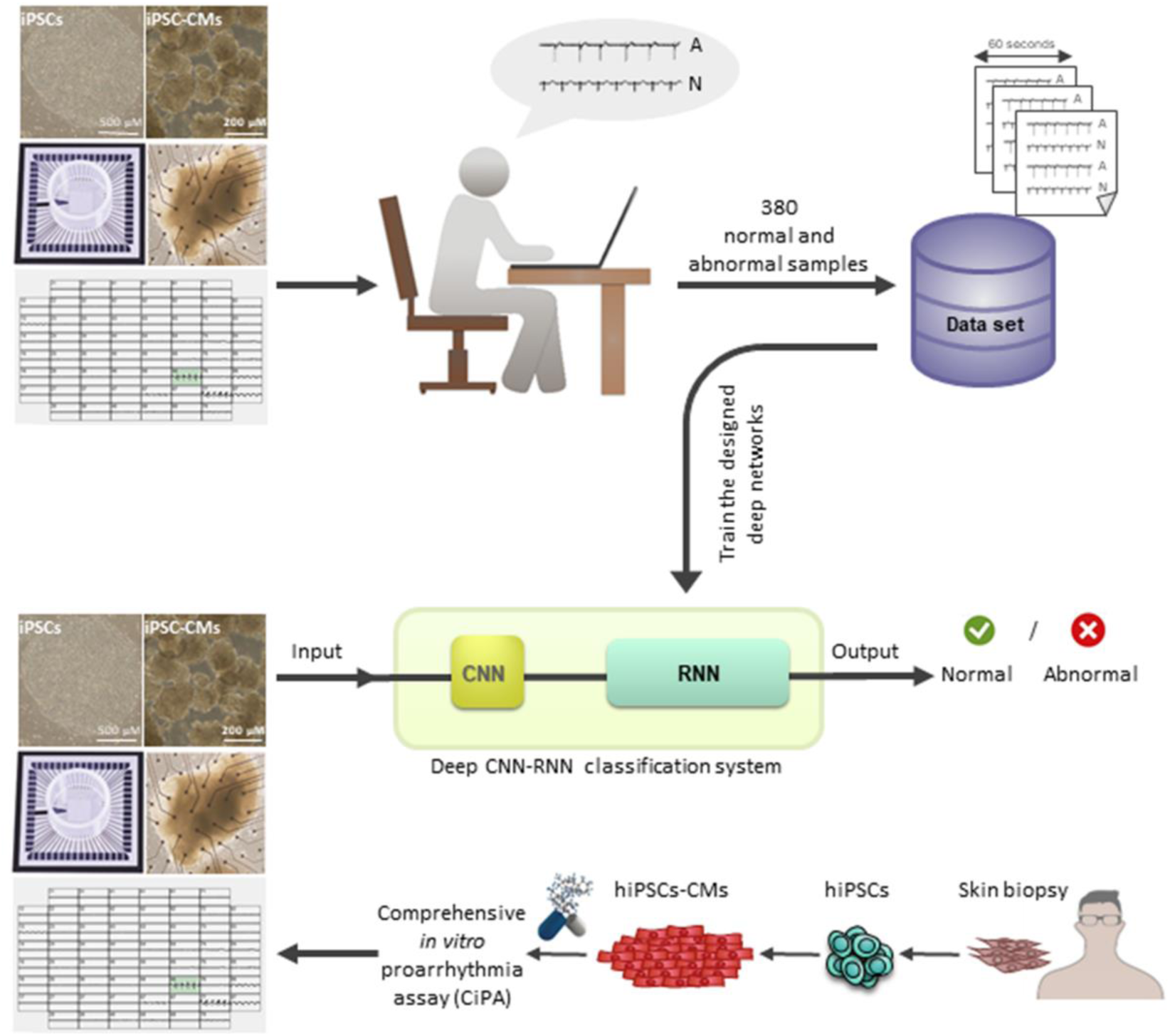
Deep learning based proarrythmia analysis for cardiac safety pharmacology. We constructed a dataset from FP recordings of iPSC-CMs using MEA system. The dataset contained 380 samples of 60-second long recordings. Each record was labeled as normal or abnormal by electrophysiology experts. Using the provided dataset, we trained the proposed deep architecture composed of CNN and RNN networks that was designed for binary classification of data. This trained deep CNN-RNN system can be further used for classification of new FP recordings into normal and abnormal categories specially in comprehensive in vitro proarrhythmia assay.

We collected a dataset of 380 field potential recordings obtained from healthy and patient-specific iPSC-CMs as well as their cardioactive drugs experiments to train the proposed classifier. These 380 recordings were labeled as normal and abnormal waveforms by electrophysiology experts. The abnormal waveforms included long FPD, bradycardia, tachycardia and polymorphic (Figure 4). We used 20% of this dataset as the test set and 80% for training and validation of the classification method. The 60-second long recordings were preprocessed and converted to small windows of 5-second long field potential recordings (Figure 1). The preprocessing resulted in about 7000 input signals. Following the training of CNN by these 5-second long recordings, the RNN was trained to aggregate output results of CNN for each waveform. The final trained networks were saved for further classification of test signals as well as any other input signal (Figure 2). The results of classification by deep learning-based classifier showed the performance metrics of 0.84 accuracy (Equation 1), 0.84 sensitivity (Equation 2), 0.85 specificity (Equation 3), and 0.88 precision (Equation 4) on the test dataset. In addition, the F-score as a harmonic mean of sensitivity and precision (Equation 5), was evaluated to be 0.86. Performance metrics were calculated by using the following formula:

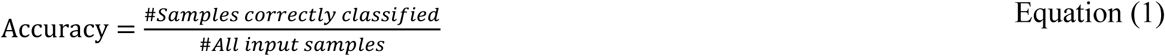

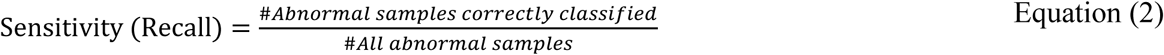

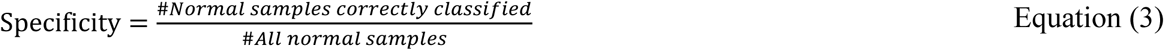

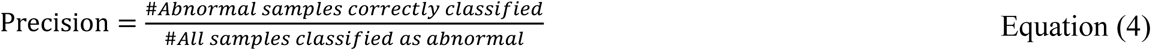

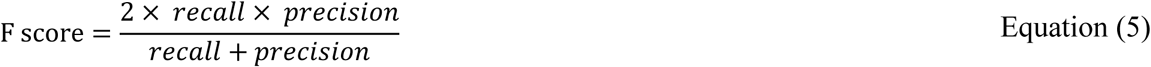

**Figure 4.**
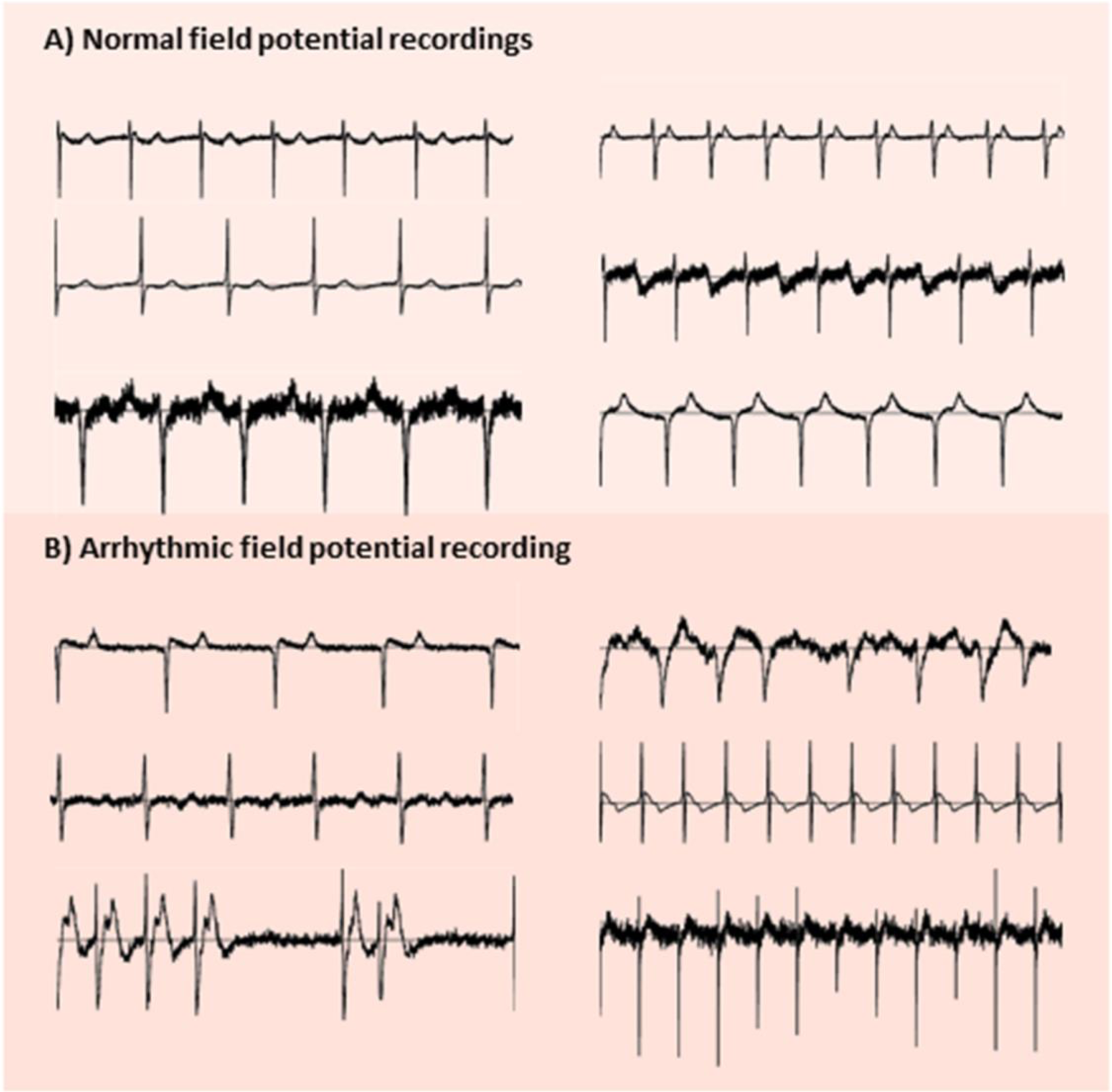
Representative extracellular field potential recordings of iPSC-CMs. A) 5-second long recordings labeled as normal. B) 5-second long traces labeled as arrhythmic due to the observed phenotypes such as long FPD, bradycardia, and polymorphic waveforms.

Therefore, the proposed classification method has resulted in a considerable performance without using the expert knowledge for designing the features and the classification system.

## Discussion

The CiPA proposes a cardiotoxicity screening schema which will result in a more robust, efficient, and sensitive system avoiding a late drug attrition from the market ^9^. The iPSC-CMs constituted a valuable tool for validating *in silico* reconstruction of pro-arrythmic risk throughout the process of drug development. Therefore, the functional assays representing the excitability or contractile function of these cells as well as advanced techniques for data analysis would be an important complement to CiPA.

To find a classifier for field potential recordings, we exploited a two-step deep learning model composed of CNN and RNN which resulted in the identification of normal and abnormal waveforms with high accuracy. Moreover, the F-score results as well as other metrics confirmed the capability of the model in classification of the recordings.

The introduction of combined iPSC-CMs technology and MEA system for drug discovery and cardiotoxicity assays has encouraged many scientists to test this platform and analyze its output signals ^32-37^. Especially, this system is developing to a high-throughput drug screening platform, highlighting the necessity for generation of automated and sensitive analysis tools. Over the last few years, the majority of analysis tools were developed for calculating the field potential parameters. Pradhapen and coworkers developed a semi-automatic software for analysis of field potentials recorded from iPSC-CMs ^15^. Their offline software called cardioMD had correlation analysis and ensemble averaging features which were used to reliably analyze the field potential durations and providing an output waveform for the expert to manually determine various morphology changing signals. In another approach, the waveforms obtained from impedance measurements of iPSC-CMs were subjected to a mathematical model (hCAR) for arrhythmic risk assessment ^14^. The hCAR model could successfully bridge the gap between hERG screening system and *in vivo* QT prolongation as well as arrhythmia assessment in preclinical and clinical studies. Furthermore, cytotoxic factors could be identified and rank-ordered using hCAR model. However, it employed some defined parameters and mathematical equations to predict proarrhythmia risk, thus limiting the power of analysis to the proposed features. In contrast, we developed a deep learning-based analysis tool by using a broad range of field potential features which resulted in arrhythmia assessment by monitoring morphological changes rather than application of any preselected features. The proposed CNN model showed the ability to process raw data directly, and did not require to use hand-designed features. Therefore, it may overcome the drawbacks of using hand-engineered features that restricts the accuracy of classification to the comprehensiveness and the quality of the designed features. In other words, CNNs have integrated the feature extraction and classification phases by learning both the representation and the classifier jointly to produce superior results. In agreement to our finding, deep CNNs have achieved spectacular results and are known as state-of-the-art methods for many tasks of medical image analysis ^20^ and biological data analysis ^38^. Furthermore, our deep learning method enables the analysis of recordings with various lengths (>5 seconds) by incorporating the designed RNN in the second stage. Today, RNN is one of the most promising methods to analyze sequential data. RNNs not only can take sequential inputs with various lengths, but also can process each element of the input sequence while maintaining a history of previous inputs in their hidden states. By using these networks, we are able to process the sequence of results obtained from the small windows as the output of the first step of our model with no limit on the length of field potential recordings. Additionally, the performance of the proposed model can be improved by increasing the size of dataset since the deep learning methods can show their real capability when they are trained on large datasets. Moreover, the binary classification can be extended to multi-class classification if larger datasets are used.

In conclusion, the proposed two-step CNN-RNN model provides an efficient, sensitive, and accurate platform for analysis and interpretation of IPSC-CM/MEAs datasets which may facilitate the preclinical cardiac safety pharmacology through CiPA.

## Acknowledgement

The authors would like to thank Dr. Maryam Barekat for her time and assistance in cardiologist-based arrhythmia detection. We are thankful to Dr. Anna Meyfour for her kind help in preparing figures.

## Supporting information

**Supplementary figure 1.** Representative field potential recordings with various morphologies.

**Supplementary figure 2.** The field potential parameters used for the analysis of recordings from iPSC-CMs.

**Supplementary movie 1.** Spontaneously beating iPSC-CMs spheroid plated on the electrodes of an MEA plate.

